# Repression of an activity-dependent autocrine insulin signal is required for sensory neuron development in *C. elegans*

**DOI:** 10.1101/481010

**Authors:** Lauren Bayer Horowitz, Julia P. Brandt, Niels Ringstad

## Abstract

Nervous system development is instructed both by genetic programs and activity-dependent refinement of gene expression and connectivity. How these mechanisms are integrated remains poorly understood. Here, we report that the regulated release of insulin-like peptides (ILPs) during development of the *C. elegans* nervous system accomplishes such an integration. We find that the p38 MAP kinase PMK-3, which is required for the differentiation of chemosensory BAG neurons, functions by limiting expression of an autocrine ILP signal that represses a chemosensory-neuron fate. ILPs are released from BAGs in an activity-dependent manner during embryonic development, and regulate neurodifferentiation through a non-canonical insulin receptor signaling pathway. The differentiation of a specialized neuron-type is, therefore, coordinately regulated by a genetic program that controls ILP expression and by neural activity, which regulates ILP release. Autocrine signals of this kind may have general and conserved functions as integrators of deterministic genetic programs with activity-dependent mechanisms during neurodevelopment.

## Introduction

The nervous system comprises many neuron-types, each endowed with a unique physiology, connectivity and molecular profile. Diversity in neuronal form and function is required for the assembly of neural circuits that support complex brain functions and behaviors. Understanding how this diversity is generated during development remains a major question in neuroscience. A remarkable feature of nervous system development is that genetically encoded developmental programs cooperate with neural activity dependent processes to promote the differentiation of specific neuron-types and instruct neuronal connectivity (Wamsley and Fishell 2017). How these two different mechanisms - one specified and the other activity-dependent - are integrated during nervous system development remains poorly understood.

The *C. elegans* nervous system displays a wide range of neuronal diversity, and is a powerful model to study neuronal differentiation (White et al. 1986; Hobert et al. 2016). The mostly invariant cell lineage that generates the *C. elegans* nervous system (Sulston 1977; Sulston 1983) suggests that neuronal differentiation in *C. elegans* is principally determined by genetic programs intrinsic to the cell-lineage. Indeed, many studies have identified transcription factors that act in specific sub-lineages to promote specific neural fates (Hobert 2016). However, there are also important roles for neuronal activity during development of the *C. elegans* nervous system. For example, there is a striking role for activity of embryonic AWC chemosensory neurons in determining their differentiation into functionally distinct subtypes (Troemel 1999; Sagasti 2001). More recently, it has been found that there is a critical period during which neural activity instructs circuit assembly in the *C. elegans* motor system (Barbagallo et al. 2017). Post-developmentally, neural and sensory activity is required for maintaining the proper morphology of chemosensory neurons (Peckol 1999; Mukhopadhyay et al. 2008), and for expression of chemosensory receptors and neuropeptides that define specific chemosensory neuron fates (Peckol et al. 2001; Gruner et al. 2014; Rojo Romanos et al. 2017). Like the vertebrate nervous system, therefore, development of the *C. elegans* nervous system requires both lineally programmed gene regulation and neural activity.

We have investigated mechanisms required for the development of a pair of *C. elegans* sensory neurons - the BAGs, which sense microbe-derived carbon dioxide (CO_2_) to control foraging behaviors (Brandt and Ringstad 2015). Properly specified BAG neurons are equipped with a chemotransduction apparatus used to sense CO_2_, which includes the receptor-type guanylyl cyclase GCY-9 (Smith et al. 2013). They also express the neurotransmitter glutamate (Serrano-Saiz et al. 2013) together with a specific set of neuropeptides *e.g.* the FMRF-amide neuropeptides FLP-17, FLP-10 and FLP-19 (Kim and Li 2004). Previous studies identified a number of transcription factors that promote the BAG neuron fate (Guillermin et al. 2011; Brandt et al. 2012; Gramstrup Petersen and Pocock 2013; Rojo Romanos et al. 2015). However mutants for these transcription factors still generate BAG neurons that differentiate to some extent. Therefore, other mechanisms that promote a BAG fate must exist. Through a screen for additional regulators of the BAG fate we identified the p38 MAP kinase (MAPK) PMK-3 (Brandt and Ringstad 2015). PMK-3 is required during development, and post-developmental expression of PMK-3 does not restore gene expression or function to *pmk-3* mutant BAG cells (Brandt and Ringstad 2015). The phenotype of *pmk-3* mutants differs from that of transcription factor mutants; *pmk-3* mutants are strongly defective in BAG-neuron function, but their gene-expression defects are restricted to a subset of BAG-specific genes. Although it is clear that PMK-3 functions in BAG development, how PMK-3 promotes differentiation of BAG neurons is unknown. p38 MAPKs have many functions in the nervous system (Thomas and Huganir 2004), but these functions are often part of injury- or stress-responses, and roles for p38 MAPKs in neuronal differentiation remain poorly understood.

To determine how PMK-3 functions in BAG neuron development, we isolated and characterized mutations that suppress the gene expression defects caused by loss of PMK-3. Here we discover that a major complementation group of suppressor mutations comprises alleles of *unc-31*, which encodes a factor required for the regulated secretion of neuropeptides and hormones. Loss of *unc-31* restores gene expression to *pmk-3* mutant BAG cells by interfering with the regulated release of insulin-like peptides (ILPs), which are overexpressed in *pmk-3* mutants and repress expression of a BAG cell fate. These ILPs are released from BAG neurons themselves and, therefore, function as an inhibitory autocrine signal during BAG neuron development. This mechanism combines a gene regulatory program, to set levels of ILP expression, and neural activity, which controls the release of ILPs during development, to regulate the differentiation of a specific neuron-type. We propose that similar mechanisms might function widely during nervous system development to integrate neural activity with genetically specified developmental programs.

## Results

### *pmk-3* mutant sensory neurons are defective in expression of a functionally important neuropeptide and also have defects in sensory transduction and synapse formation

*pmk-3* mutants fail to express the BAG-neuron-specific FMRF-amide neuropeptide *flp-17* (**Figure 1A**). FLP-17 peptides activate the G_i/o_-coupled receptor EGL-6 to inhibit motor neurons in the *C. elegans* egg laying system (Ringstad and Horvitz 2008), and were recently shown to also be required for BAG-neuron-dependent CO_2_ avoidance behavior (Guillermin et al. 2017; Lee et al. 2017). To determine whether the behavioral defect of *pmk-3* mutants can be explained by their failure to express *flp-17* neuropeptides, we compared the CO_2_ avoidance defects of *pmk-3* and *flp-17* mutants. Both *pmk-3* and *flp-17* mutants were severely defective for CO_2_ avoidance, although *pmk-3* mutants displayed a more severe defect (**Figure 1B**). Interestingly, we found that *egl-6* mutants, which lack the known receptor for FLP-17 peptide, were wild-type for CO_2_ avoidance (**Figure 1B**), suggesting that FLP-17 acts on distinct receptors in circuits that mediate chemotaxis.

**Figure 1:**
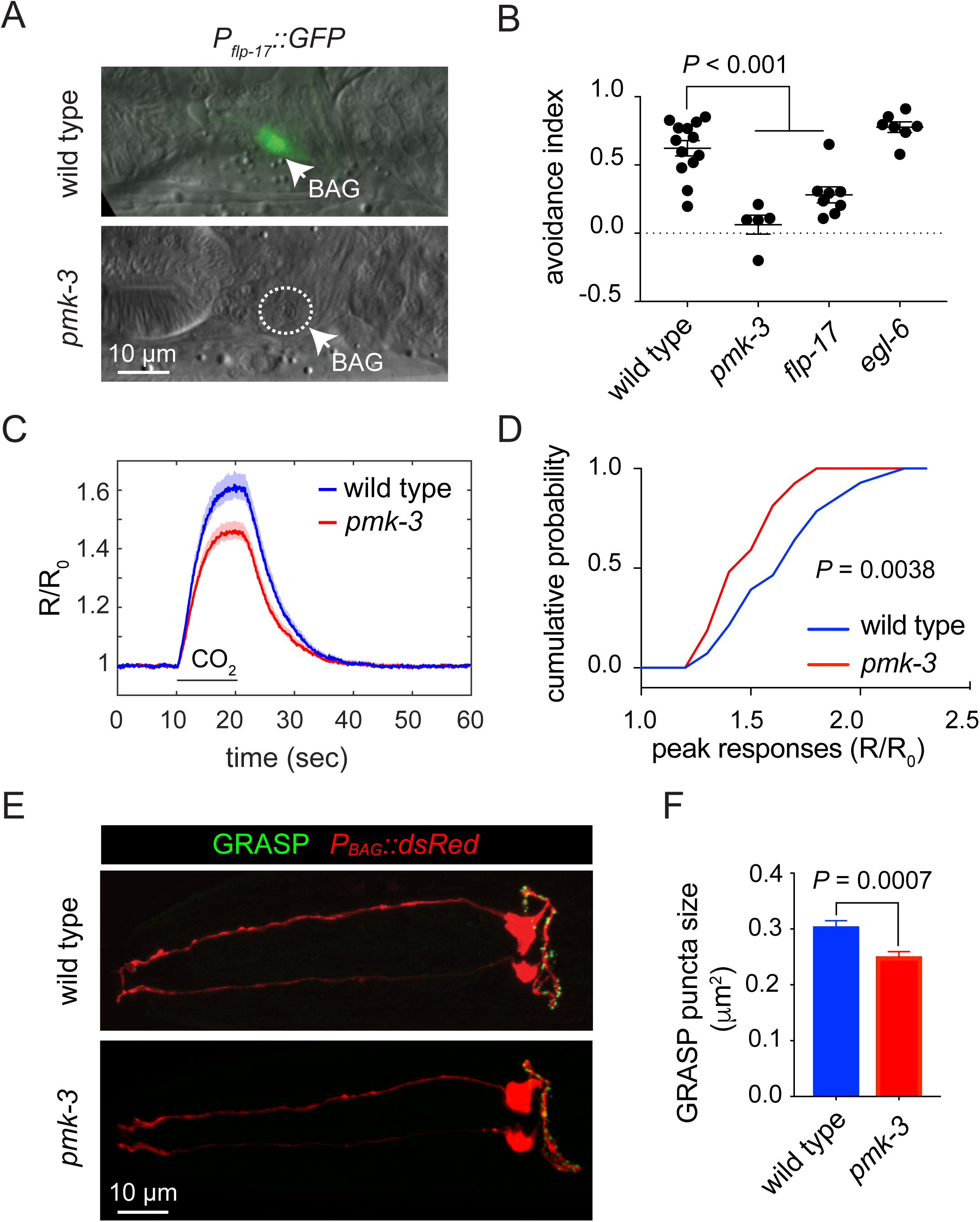
PMK-3 mutation affects neuropeptide expression, chemotransduction and synapse morphology in chemosensory BAG neurons. A. Overlaid differential interference contract (DIC) and fluorescence micrographs of wild-type and *pmk-3(wz31)* mutant animals expressing *P*_*flp-17*_*∷GFP*, a marker of differentiated BAG cells. B. CO_2_-avoidance indices of the wild type, *pmk-3(ok169), flp-17(n4894)*, and *egl-6(n4536)* mutants. N >= 5 independent trials. *P* < 0.0001 for wild type vs. *pmk-3*, and *P* = 0.0007 for wild-type vs. *flp-17*, ordinary one-way ANOVA followed by Tukey’s multiple comparisons test. Error bars represent SEM. C. Average calcium responses of wild-type and *pmk-3(ok169)* mutant BAG cells to 10% CO_2_ stimuli. Shaded area represents the mean ± SEM. N > 26 animals/genotype. (R/R_0_) is the ratio of YFP/CFP emissions normalized to the pre-stimulus ratio. D. Cumulative probability plots of the peak calcium responses (R/R_0_) in wild-type and *pmk-3(ok169)* mutant BAG cells to CO_2_ stimuli. *P* = 0.0038, unpaired t-test. E. Fluorescence micrographs of BAG GRASP puncta in wild-type and *pmk-3(ok169)* mutant BAG cells using *P*_*gcy-9*_*∷nlg-1∷GFP*_*1-10*_ *P*_*odr-2b*_*∷nlg-1∷GFP*_*11*_, overlaid with *P*_*gcy-9*_*∷dsRed* to label BAG cells. F. Quantification of the size of GRASP puncta in wild-type and *pmk-3(ok169)* mutant BAG cells. N > 28 animals/genotype. *P* = 0.0007, unpaired t-test. Error bars represent SEM.

Because *pmk-3* mutants exhibited a stronger CO_2_ avoidance defect than *flp-17* mutants, we sought to determine whether PMK-3 regulates other aspects of BAG-neuron physiology or structure. We first tested whether *pmk-3* mutant BAG cells exhibit sensory transduction defects using *in vivo* calcium imaging to measure their calcium responses to CO_2_ stimuli (Brandt et al. 2012). *pmk-3* mutant BAG cells responded to strong CO_2_ stimuli, but their responses were significantly smaller than those of the wild type (**Figure 1C-D, Figure S1**). When wild-type and *pmk-3* mutant cell responses were scaled to unity, we observed no difference between the dynamics of the calcium responses, indicating that the kinetics of cell activation was not altered by *pmk-3* mutation (**Figure S1**).

We next tested whether *pmk-3* mutant BAG cells form proper synapses using GFP-Reconstitution across Synaptic Partners (GRASP) (Feinberg et al. 2008) to label a subset of BAG synapses. GRASP-puncta were observed throughout the axons of both wild type and *pmk-3* mutant BAG cells (**Figure 1E**). We observed no significant difference in the number of puncta in *pmk-3* mutants and in the wild type (**Figure S2**). The average size of puncta in *pmk-3* mutants, however, was 18% smaller than that of the wild type (**Figure 1F**). From these data we concluded that the behavioral defects of *pmk-3* mutants that are caused by their failure to express FLP-17 neuropeptides are likely exacerbated by accompanying defects in BAG neuron chemotransduction and synapse formation.

### Genes required for Ca^2+^-dependent neural secretion suppress a neuropeptide gene expression defect of *pmk-3* mutants

Previously, we found evidence that PMK-3 acts downstream of the Toll-like Receptor TOL-1 to promote BAG neuron development (Brandt and Ringstad 2015). How PMK-3 itself regulates expression of a BAG neuron fate was unknown. To address this question, we performed a genetic screen for suppressors of the *flp-17* expression defects of *pmk-3* mutants. From two screens that covered approximately 25,000 mutagenized haploid genomes, we recovered 18 mutants in which BAG neuron expression of *flp-17* was restored to *pmk-3* mutants (**Figure 2A-B**). Five of these mutants defined a major complementation group, and we decided to further characterize the affected gene. These suppressor mutations strongly restored the penetrance of *flp-17* reporter expression in *pmk-3* mutant BAG cells (**Figure 2C**), and also significantly restored the levels of reporter expression (**Figure S3**). We asked whether the suppressor mutations also affected another PMK-3 regulated gene. Expression of *gcy-33*, which encodes a guanyly cyclase required for sensing hypoxia (Zimmer et al. 2009) and is regulated by PMK-3 in BAG (Brandt and Ringstad 2015), was not restored in suppressed *pmk-3* mutants (**Figure S4**). Therefore the suppressor gene defined by this complementation group has strong effects on *flp-17* expression, but does not completely restore the *pmk-3* phenotype to wild-type.

**Figure 2:**
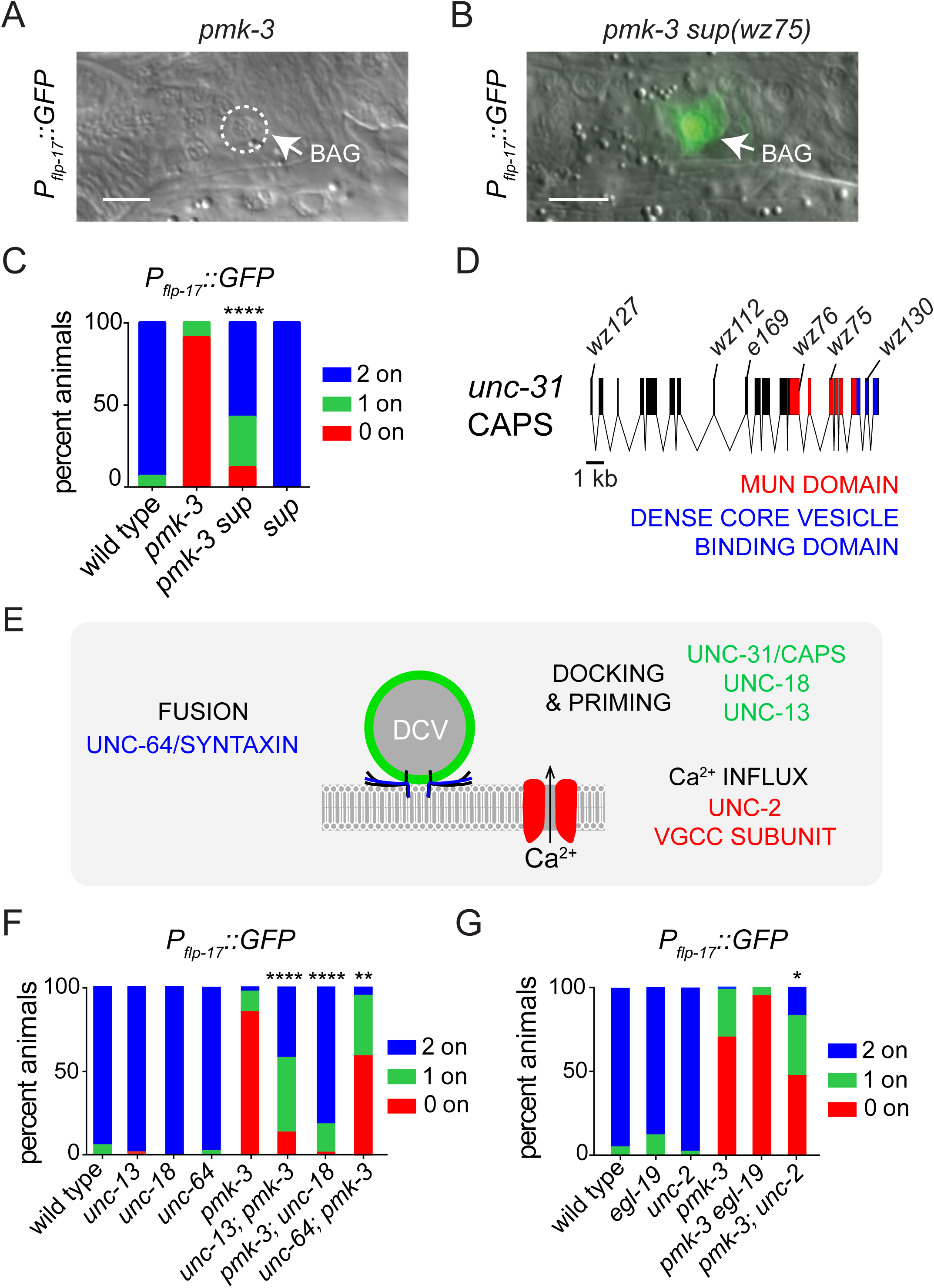
Mutation of genes required for regulated neuronal secretion suppress the defects caused by loss of PMK-3. A-B. Overlays of DIC and fluorescence micrographs of a *pmk-3(wz31)* mutant (A) and a *pmk-3(wz31)* mutant animal carrying the suppressor mutation *wz75* (B) expressing *P*_*flp-17*_*∷GFP*. (Scale bar:10 μm). C. Penetrance of *P*_*flp-17*_*∷GFP* expression in the wild type, *pmk-3(wz31), pmk-3(wz31) sup(wz75)* mutant animals, and in animals carrying a mutation in the suppressor gene on its own, *sup(e169)*. D. Structure of the *unc-31* locus. *wz75, wz76, wz112, wz127* and *wz130* are 5 non-complementing alleles isolated by our *pmk-3* suppressor screen, and *e169* is the *unc-31* reference allele. E. Genes that function with UNC-31/CAPS to regulate secretion of dense core vesicles (DCVs). (VGCC) Voltage-Gated Calcium Channel. F. Penetrance of *P*_*flp-17*_*∷GFP* expression in the wild type, *unc-13(e51), unc-18(e81), unc-64(e246), pmk-3, unc-13(e51); pmk-3(ok169), pmk-3(ok169); unc-18(e81)*, and *unc-64(e246); pmk-3(wz31)* mutants. Data shown for *pmk-3* represent the average penetrance of *P*_*flp-17*_*∷GFP* expression collected from *pmk-3(wz31)* and *pmk-3(ok169)* mutant animals. G. Penetrance of *P*_*flp-17*_*∷GFP* expression in the wild type, *egl-19(n582), unc-2(e55), pmk-3(ok169), pmk-3(ok169) egl-19(n582)*, and *pmk-3(ok169); unc-2(e55)* mutants. N > 25 animals/ genotype. (*) *P* < 0.05; (**) *P* < 0.01, (****) *P* < 0.0001, chi-square test.

Using high-resolution Single Nucleotide Polymorphism (SNP) mapping and whole-genome sequencing we identified the suppressor gene as *unc-31* (**Figure 2D**), which encodes the *C. elegans* homolog of <u>C</u>alcium-dependent <u>A</u>ctivator <u>P</u>rotein for <u>S</u>ecretion (CAPS). UNC-31/CAPS is a neuron-specific factor required for docking and priming of dense core vesicles (DCVs), which mediate the regulated release of neuropeptides, hormones and growth factors (Speese et al. 2007; Zhou et al. 2007). Because of the known function of *unc-31*, we hypothesized that Ca^2+^-dependent secretion plays a role in BAG neuron development. We tested whether mutations that affect other critical factors for neural secretion (**Figure 2E**) also suppress *pmk-3*. Mutation of either UNC-13 or UNC-18, which like UNC-31 are required for docking and priming of synaptic and dense core vesicles (Richmond 1999; Weimer et al. 2003), strongly restored the frequency of *flp-17* expression to *pmk-3* mutants (**Figure 2F**). We also tested a partial loss-of-function allele of the syntaxin homolog, UNC-64 (Saifee 1998), and observed partial restoration of *flp-17* expression to *pmk-3* mutant BAG neurons (**Figure 2F**).

Because DCV exocytosis is triggered by elevated Ca^2+^ that enters cells through voltage-gated calcium channels (VGCCs), we next tested whether mutation of VGCC subunits suppresses the gene expression defect of *pmk-3* mutants. Mutation of the L-type VGCC alpha-1 subunit, *egl-19* (Lee 1997), did not restore *flp-17* expression to *pmk-3* mutants (**Figure 2G**). By contrast, mutations in the NPQ-type VGCC alpha-1 subunit, *unc-2* (Schafer 1995), did suppress *pmk-3* and significantly restored *flp-17* expression (**Figure 2G, Figure S5B**). The VGCC alpha-2-delta subunit UNC-36 is a part of both L- and NPQ-type channels (Lee 1997). Partial loss of UNC-36 function did not affect the frequency with which *pmk-3* mutants expressed *flp-17* but did cause an increase in the expression levels of *flp-17* when it was expressed in *pmk-3* mutants (**Figure S5**). Together, these data show that Ca^2+^-dependent secretion and neural activity antagonize PMK-3 dependent neuropeptide expression in BAG neurons.

### Neural activity during development regulates neuropeptide gene expression in BAG neurons

*pmk-*3 functions cell-autonomously and during a critical period of embryonic development to promote expression of *flp-17* in BAG neurons (Brandt and Ringstad 2015). We tested whether neural activity and regulated secretion also act cell-autonomously in BAG neurons during the same critical period. To dampen BAG activity, we used the BAG-neuron-specific and PMK-3-independent *gcy-9* promoter (Smith et al. 2013; Brandt and Ringstad 2015) to overexpress the inward rectifying potassium (K^+^) channel IRK-1 (Emtage et al. 2012). IRK-1 expression in BAGs restored *flp-17* expression to *pmk-3* mutants (**Figure 3A**). In parallel, we performed BAG-targeted knockdown of *unc-31* using RNAi. Knock-down of *unc-31* in BAGs alone restored *flp-17* expression to *pmk-3* mutant BAG neurons (**Figure 3B**). We compared the effects of BAG-targeted IRK-1 expression or *unc-31* knock-down to the effects of disrupting neural activity and regulated secretion in all neurons using a pan-neuronal promoter. BAG-targeted inhibition of regulated secretion or neural activity suppressed *pmk-3* as strongly as the corresponding pan-neuronal manipulations (**Figure 3A,B**). These data strongly suggest that neural activity and UNC-31 function cell autonomously in BAGs to regulate *flp-17* expression.

**Figure 3:**
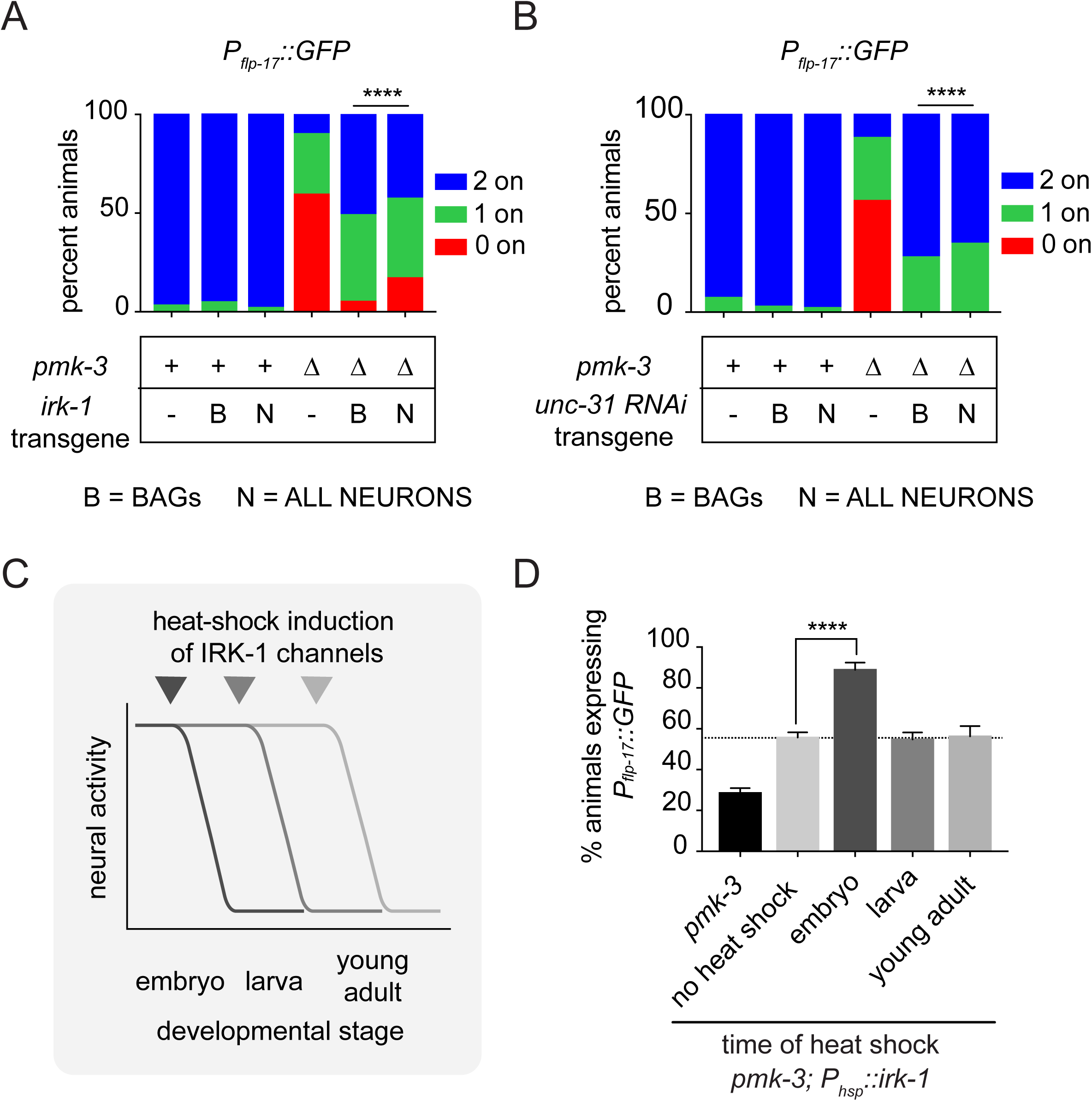
Electrical activity in BAG neurons during development regulates expression of a PMK-3-regulated gene. A. Silencing BAG neural activity using *P*_*gcy-9*_*∷irk-1* strongly restores the penetrance of *P*_*flp-17*_*∷GFP* expression to *pmk-3(ok169)* mutant BAG cells. Similar effects are observed when pan-neuronal activity is silenced using *P*_*rab-3*_*∷irk-1*. B. BAG-targeted knockdown of *unc-31* using RNAi, *P*_*gcy-9*_*∷unc-31 RNAi*, is sufficient to strongly restore the penetrance of *P*_*flp-17*_*∷GFP* expression to *pmk-3(ok169)* mutant BAG cells, similar to pan-neuronal RNAi of *unc-31, P*_*rab-3*_*∷unc-31 RNAi*. N ≥ 25 animals/genotype. (****) *P* < 0.0001, chi-square test. C. Animals carrying a transgene for heat-shock-inducible overexpression of *irk-1, P*_*hsp*_*∷irk-1*, were heat-shocked at different developmental stages (embryonic, larval or adult) to induce *irk-1* expression and reduce neural activity. D. Percentage of animals expressing *P*_*flp-17*_*∷GFP* in *pmk-3(ok169)* and *pmk-3(ok169); P*_*hsp*_*∷irk-1* mutant animals that had been heat shocked at the indicated developmental stages. N > 95 animals for all measurements. (****) *P* < 0.0001, ordinary one-way ANOVA followed by Dunnet’s multiple comparison test. Bar graph data are plotted as means ± SEM.

We next tested when neural activity is required to exert its effect on *flp-17* expression. We expressed *irk-1* under the control of a heat-shock-inducible promoter in *pmk-3* mutants. Animals were heat-shocked as embryos, larvae, or adults to transiently induce *irk-1* expression and reduce neural activity (**Figure 3C**), and were then scored for *flp-17* expression as adults. The transgene for inducible expression of IRK-1 conferred some suppression in the absence of heat shock, suggesting that it supports some basal expression of IRK-1 (**Figure 3D**). However, heat-shock of embryos restored *flp-17* expression to *pmk-3* mutants above this baseline (**Figure 3D**). By contrast, induction of IRK-1 expression during larval or adult stages did not significantly modify the *pmk-3* gene expression defect (**Figure 3D**). Together, these data indicate that, like PMK-3, neural activity and UNC-31 are required during development in BAG neurons to regulate neuropeptide gene expression.

### *pmk-3* mutation dysregulates expression of insulin-like genes in BAG neurons

Why might mutations in *unc-31* and other genes required for regulated secretion of peptides suppress the effects of *pmk-3* mutation? We hypothesized that in *pmk-3* mutants a factor, possibly a peptide hormone, is secreted by BAGs in an UNC-31-dependent manner and inhibits their development. We sought evidence for such a secreted factor by transcriptionally profiling embryonic BAG neurons from the wild type and *pmk-3* mutants. We used fluorescence-activated cell-sorting to purify GFP-marked BAG neurons and ‘non-BAG’ non-fluorescent cells. Differential gene expression analysis identified 692 transcripts whose expression significantly differed between wild-type and *pmk-3* mutant BAG cells (*P* < 0.05) (**Figure 4A**). 189 of these transcripts were also BAG-enriched, as determined by a comparison of wild-type BAG neurons vs non-BAG cells (*P* < 0.05). Principal component analysis revealed that wild-type and *pmk-3* mutant BAG cell transcriptomes are highly separable (**Figure 4B**), indicating that *pmk-3* is required for proper expression of many genes in developing BAG neurons.

**Figure 4:**
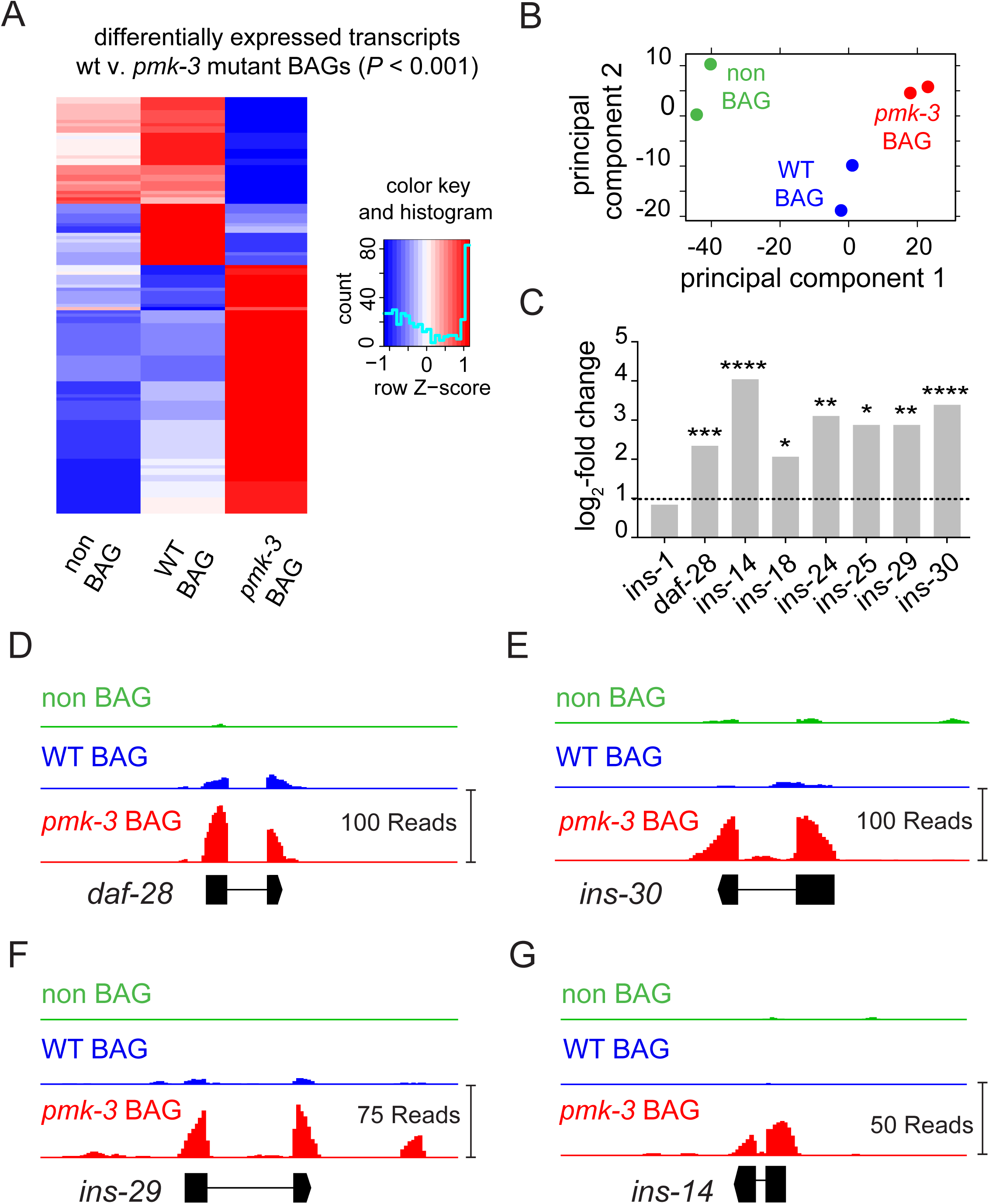
PMK-3 represses expression of genes encoding insulin-like peptides in BAG chemosensory neurons. A. Heatmap showing the relative expression of the most differentially expressed genes (*P* < 0.001, 129 transcripts) between wild-type (WT) and *pmk-3(wz31)*. Expression of these transcripts in non-BAG cells is also shown. Colors represents standardized Z-scores calculated from the average normalized DeSeq2 read counts, with blue representing low expression and red representing high expression. B. Principal component analysis of the transcriptomes from wild-type BAG cells, *pmk-3(wz31)* mutant BAG cells, and non-BAG cells. Each dot represents one biological replicate. C. Fold-changes of gene expression for insulin like peptides (ILPs) in *pmk-3(wz31)* mutant BAG cells versus wild-type BAG cells. N = 2 biological replicates. (*) *P* < 0.05. (**) *P* < 0.01. (***) *P* < 0.001. (****) *P* < 0.0001. *P* values were adjusted for False Discovery Rates (FDR) using DeSeq2 (Love et al. 2014). (D-G). Read coverage histograms for a subset of indicated ILPs that are significantly overexpressed in *pmk-3(wz31)* mutant BAG cells. (see Materials and Methods).

Inspection of genes affected by *pmk-3* mutation revealed that *pmk-3* mutant BAG cells over-express multiple insulin-like peptides (ILPs) (**Figure 4C**). The ILP gene *daf-28*, which is enriched in wild-type BAG neurons compared to non-BAG cells (**Figure S6**), is upregulated in *pmk-3* mutant BAGs (**Figure 4D**). *ins-14, ins-29* and *ins-30*, which are not normally expressed in wild-type BAGs, are clearly expressed in *pmk-3* mutant BAG neurons (**Figure 4E-G**). The ILPs affected by *pmk-3* mutation all encode ILPs that are agonists of the *C. elegans* insulin/IGF-like receptor (InR) DAF-2 (Kenyon 1993; Kimura 1997; Murphy and Hu 2013). Our RNA-Seq data also revealed that an ILP gene that encodes an InR antagonist -*ins-1* (Cornils et al. 2011) - is enriched in wild-type BAG neurons (**Figure S6**), but its expression is not affected by *pmk-3* mutation (**Figure 4C**). These data suggested that defects caused by loss of PMK-3 might result from excess expression of some ILPs expressed in embryonic BAGs. To test this hypothesis, we next sought to determine how ILP production by BAGs regulates BAG neuron development.

### An autocrine insulin signaling pathway antagonizes PMK-3 dependent gene expression in BAG cells

To disrupt ILP production we used the dominant negative *daf-28(sa191)* allele (**Figure 5A**). *daf-28(sa191)* generates a non-functional ILP that mis-folds and disrupts production of other ILPs (Li et al. 2003). *daf-28(sa191)* phenocopied mutations that affect regulated secretion and restored *flp-17* expression to *pmk-3* mutants (**Figure 5B**). A deletion allele of *daf-28* (**Figure 5A**) also partially restored *flp-17* expression to *pmk-3* (**Figure 5C**), indicating that although BAGs produce several ILPS, DAF-28 has a non-redundant function in regulating BAG gene expression. We next overexpressed DAF-28(sa191) specifically in *pmk-3* mutant BAG neurons to test whether ILP production in BAG neurons themselves is required for their development. Overexpressing DAF-28(sa191) in BAGs strongly restored *flp-17* expression (**Figure 5B**). Together, these data indicate that ILP expression by BAGs antagonizes PMK-3-dependent expression of *flp-17*. Because *pmk-3* mutant BAGs overexpress ILPs, these data further suggest that the BAG neuron defects of *pmk-3* mutants are at least partly caused by dysregulated expression and release of ILPs.

**Figure 5:**
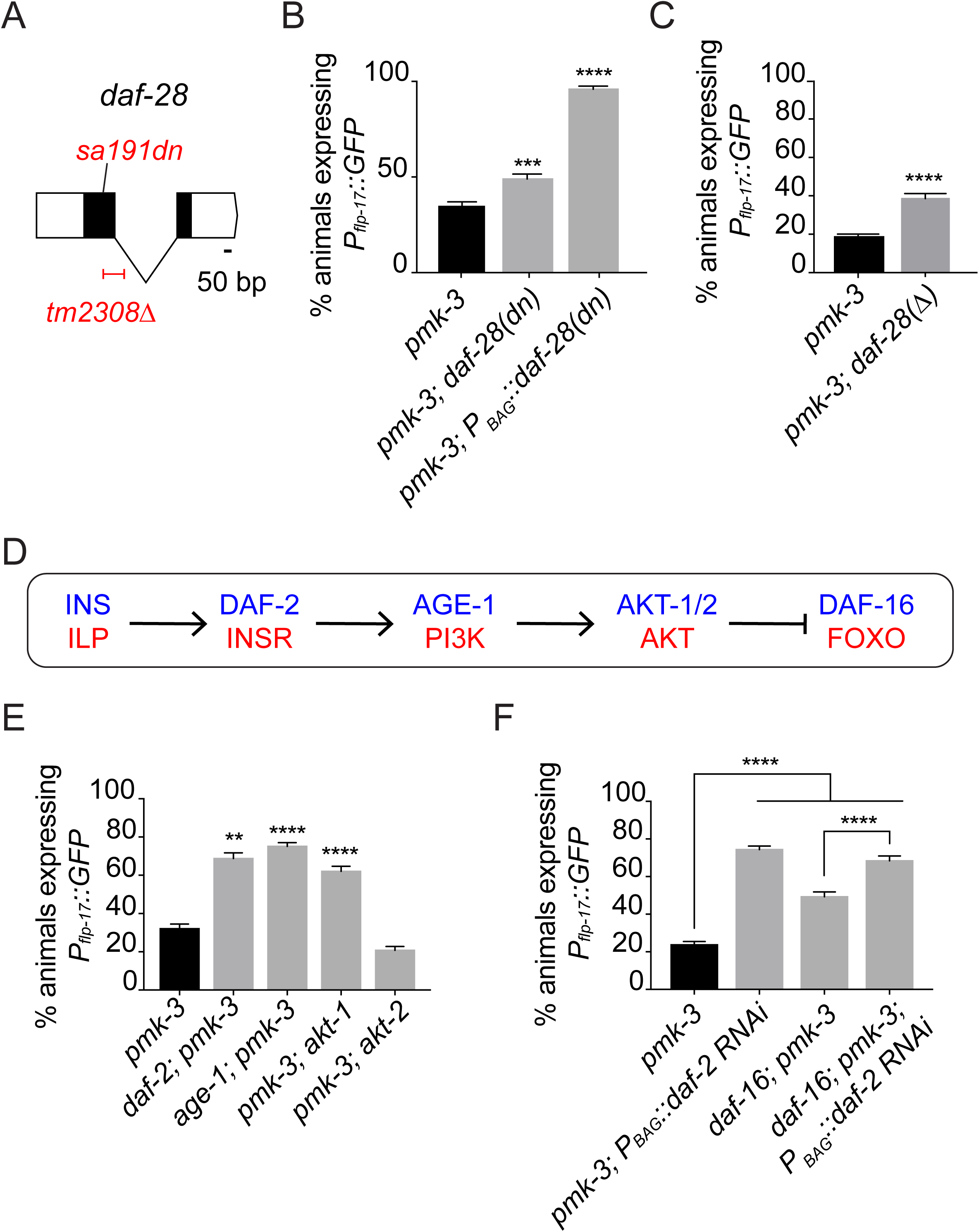
Autocrine insulin signaling antagonizes PMK-3*-*dependent neuropeptide expression in BAG neurons. A. Structure of the *daf-28* locus showing the null allele, *tm2308Δ*, and the dominant negative (dn) *sa191* allele, a point mutation that disrupts production of multiple insulin-like peptides. B. Disrupting insulin production in *pmk-3(ok169)* mutant BAG cells by overexpressing *daf-28(sa191), P*_*BAG*_*∷daf-28(sa191)*, strongly restores *P*_*flp-17*_*∷GFP* expression to *pmk-3(ok169)* mutants. The chromosomal *daf-28(sa191)* mutation also partially restores the percentage of *pmk-3(ok169)* mutants expressing *P*_*flp-17*_*∷GFP*. (***) *P* = 0.0002, (****) *P* < 0.0001, unpaired t-test. C. *daf-28(tm2308)* partially restores the percentage of *pmk-3(ok169)* mutant animals expressing *P*_*flp-17*_*∷GFP*. (****) *P* < 0.0001, unpaired t-test. D. A canonical insulin signaling pathway conserved between nematodes and vertebrates. E. Percent animals expressing *P*_*flp-17*_*∷GFP* in *pmk-3(ok169), daf-2(e1370); pmk-3(ok169), age-1(hx546); pmk-3(ok169), pmk-3(ok169); akt-1(ok525)*, and *pmk-3(ok169); akt-2(ok393)* mutant animals. For the *pmk-3* versus *daf-2; pmk-3* and *pmk-3* versus *age-1; pmk-3* comparisons, *P* = 0.0016 and *P* < 0.0001, respectively, unpaired t-test. For the *pmk-3* versus *pmk-3; akt-1* and *pmk-3* versus *pmk-3; akt-2* comparisons, *P* < 0.0001 and *P* = 0.9524, respectively, ordinary one-way ANOVA followed by Dunnet’s multiple comparisons test. F. BAG cell-targeted knockdown of *daf-2* using RNAi is sufficient to strongly restore *P*_*flp-17*_*∷GFP* expression to *pmk-3(ok169)* mutant BAG cells, even in the absence of *daf-16(wz151)*. (****) *P* < 0.0001. For *pmk-3; P*_*BAG*_*∷daf-2 RNAi* versus *daf-16; pmk-3; P*_*BAG*_*∷daf-2 RNAi, P* = 0.2612. *P* – values were calculated with an ordinary one-way ANOVA followed by Tukey’s multiple comparisons test. N >= 45 animals/genotype. Bar graph data are plotted as means ± SEM.

ILPs released from BAGs might act on other cells that in turn release factors that regulate gene expression in BAGs. Alternatively, ILPs released from BAGs might function as autocrine signals and activate insulin-receptor signaling pathways in BAG themselves. To resolve these possibilities, we sought to determine the site of action of the InR signaling pathway that regulates *flp-17* expression in BAG neurons. First, we interrogated genes known to function in InR signaling for suppression of *pmk-3*. The InR DAF-2 signals via the phosphoinositide-3 (PI-3) kinase, AGE-1 (Morris 1996), and two serine/threonine kinases, AKT-1 and AKT-2 (Paradis 1998) (**Figure 5D**). Many genes that function in the canonical InR signaling pathway mutate to suppress *pmk-*3; loss-of-function mutations in *daf-2, age-1*, and *akt-1*, each strongly restored *flp-17* expression to *pmk-3* mutants (**Figure 5E**). We observed no effect, however, of *akt-2* mutation on the *pmk-3* phenotype (**Figure 5E**). We next determined where the DAF-2 InR was required for its role in the BAG neurons development. We tested the hypothesis that ILPs are part of an autocrine signal, and performed BAG neuron-targeted knockdown of *daf-2* using RNAi. Like the *daf-2* chromosomal mutation, BAG-targeted *daf-2* RNAi strongly restored *flp-17* expression to *pmk-3* mutant BAG cells (**Figure 5F**). Together with our analysis of ILP expression in BAGs, these data indicate that an autocrine ILP signal represses *flp-17* expression in BAG neurons.

### *pmk-3* mutant BAG cells experience increased insulin signaling

InR-dependent phosphorylation inhibits entry of the Forkhead (FOXO) transcription factor DAF-16 into the nucleus to regulate transcription (Lee 2001; Lin 2001). We used this phenomenon to independently test the hypothesis that *pmk-3* mutation causes BAG neurons to experience increased autocrine ILP signaling. We generated transgenic animals that express a DAF-16∷GFP fusion in BAG neurons, and we measured the ratio of nuclear DAF-16 to cytoplasmic DAF-16 as an index of InR signaling in BAGs. This ratio varied among wild-type BAG neurons, some of which showed little nuclear DAF-16∷GFP fluorescence and others with nuclear fluorescence that was comparable to that in the cytoplasm (**Figure 6A, left, middle**). While *pmk-3* mutant BAG neurons also displayed a range of nuclear-to-cytoplasmic DAF-16∷GFP ratios, their distribution was significantly shifted towards lower ratios compared to those measured in the wild type (**Figure 6B**), and on average had more cytoplasmic DAF-16∷GFP. As expected, mutation of the InR DAF-2 caused a dramatic accumulation of DAF-16∷GFP in the nuclei of BAG neurons (**Figure 6A, right**), but in *daf-2* mutants there was no significant effect of *pmk-3* mutation on the ratios of nuclear to cytoplasmic DAF-16∷GFP (**Figure 6C**). These data indicate that *pmk-3* mutant BAG neurons experience elevated insulin signaling, and provide independent corroboration of a model in which PMK-3 negatively regulates ILP expression and InR signaling.

**Figure 6:**
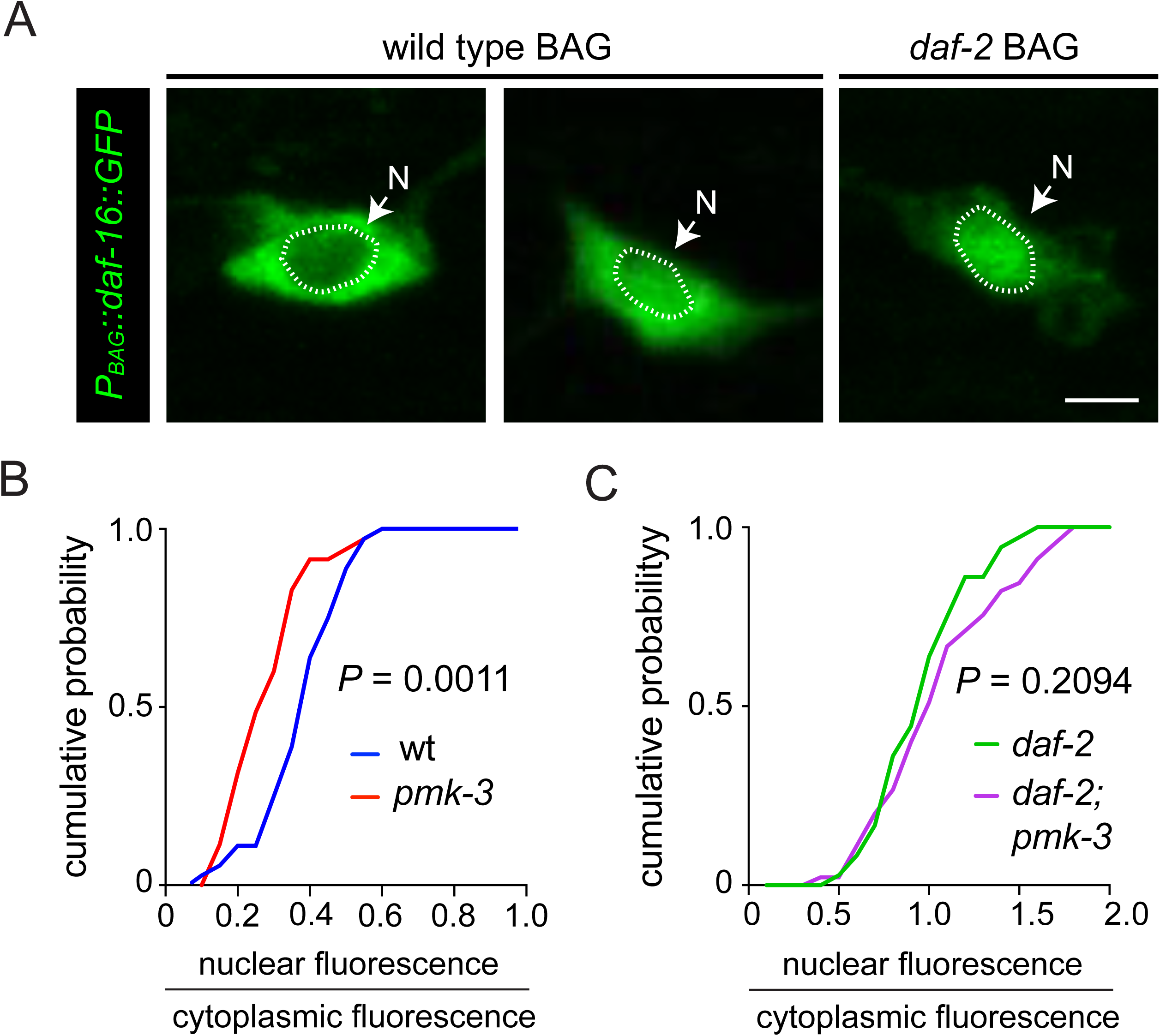
*pmk-3* mutant BAG neurons experience increased insulin signaling. A. Fluorescence micrographs of wild-type and *daf-2(e1370)* mutant animals expressing GFP-tagged DAF-16 in the BAG cell, *P*_*BAG*_*∷daf-16∷GFP*. (Scale bar: 5 μm). (N) nucleus. B. Cumulative probability plots of the nuclear to cytoplasmic DAF-16∷GFP fluorescence ratios in wild-type and *pmk-3(ok169)* mutant BAG cells. N = 36 cells from 27 animals for the wild-type and N = 35 cells from 23 animals for *pmk-3* mutants. *P* = 0.0011, unpaired t-test. C. The nuclear-to-cytoplasmic DAF-16∷GFP fluorescence ratio is not significantly different between *daf-2(e1370)* and *daf-2(e1370); pmk-3(ok169)* mutant animals. N = 46 cells from 30 animals for *daf-2* mutants and N = 45 cells from 28 animals for *daf-2; pmk-3* mutant animals. *P* = 0.2094, unpaired t-test.

### InR/DAF-2 regulates neuropeptide expression independent of FOXO/DAF-16 in BAG chemosensory neurons

We next tested whether the DAF-16/FOXO transcription factor, which is a canonical effector of InR signaling, is required for ILPs to regulate *flp-17* expression. Because the *daf-16* locus is tightly linked to the *flp-17* reporter that we use to assay the *pmk-3* mutant phenotype, we used CRISPR/Cas9 to generate a new *daf-16* deletion allele in a strain carrying the *flp-17* reporter (**Figure S7**). We did not observe an effect of *daf-16* mutation on the frequency of animals that express *flp-17*, but we did note that these mutants express *flp-17* at lower levels compared to the wild-type (**Figure S7**). Notably, knock-down of *daf-2* by RNAi still restored expression of *flp-17* to *pmk-3* mutant BAG cells in the absence of *daf-16* (**Figure 5F**), indicating that DAF-2 regulates *flp-17* expression in the absence of DAF-16. Mutating another transcription factor that functions in the DAF-2 signaling pathway – the Nrf-like transcription factor SKN-1 (Tullet et al. 2008) – did not affect the penetrance or levels of *flp-17* expression (**Figure S7**). Together, these data indicate that DAF-2 regulates gene expression in BAG neurons through a mechanism independent from its canonical effector DAF-16. Although we could not rule out a role for SKN-1 in this pathway, the absence of any effect of *skn-1* mutation on *flp-17* expression suggests that this factor does not function in BAG neuron development.

### Attenuating autocrine insulin signaling restores function to *pmk-3* mutant BAG neurons

Mutation of PMK-3 dysregulates expression of ILP genes in BAG neurons, but also affects expression of many other genes (**Figure 4A**). To what extent is BAG neuron function affected by increased production of ILPs? To address this question we tested whether disrupting autocrine insulin signaling in BAGs restores CO_2_ avoidance behavior to *pmk-3* mutants, which overexpress ILPs. We again used the dominant negative DAF-28(sa191) variant to disrupt insulin production in BAG neurons of *pmk-3* mutants, this time testing for effects of knocking down ILP production on behavior. We found that the chromosomal *daf-28(sa191)* mutation significantly restored CO_2_ avoidance behavior to *pmk-3* mutants (**Figure 7A**). Overexpressing DAF-28(sa191) in *pmk-3* mutant BAG cells also suppressed the CO_2_ avoidance defect of *pmk-3* mutants (**Figure 7B**), indicating that ILP production by BAGs is required for the effects of *pmk-3* mutation on behavior. Next, we asked whether disrupting *daf-2* function in BAGs could also restore behavior to *pmk-3* mutants. BAG-neuron-specific *daf-2* RNAi did not affect CO_2_ avoidance by wild-type worms (**Figure 7C**), but significantly restored CO_2_ avoidance to *pmk-3* mutants (**Figure 7C**). Together these data show that the functional defects caused by loss of PMK-3, which are associated with widespread changes in gene regulation, can be rescued by targeting the ILP signaling pathway in BAG neurons.

**Figure 7:**
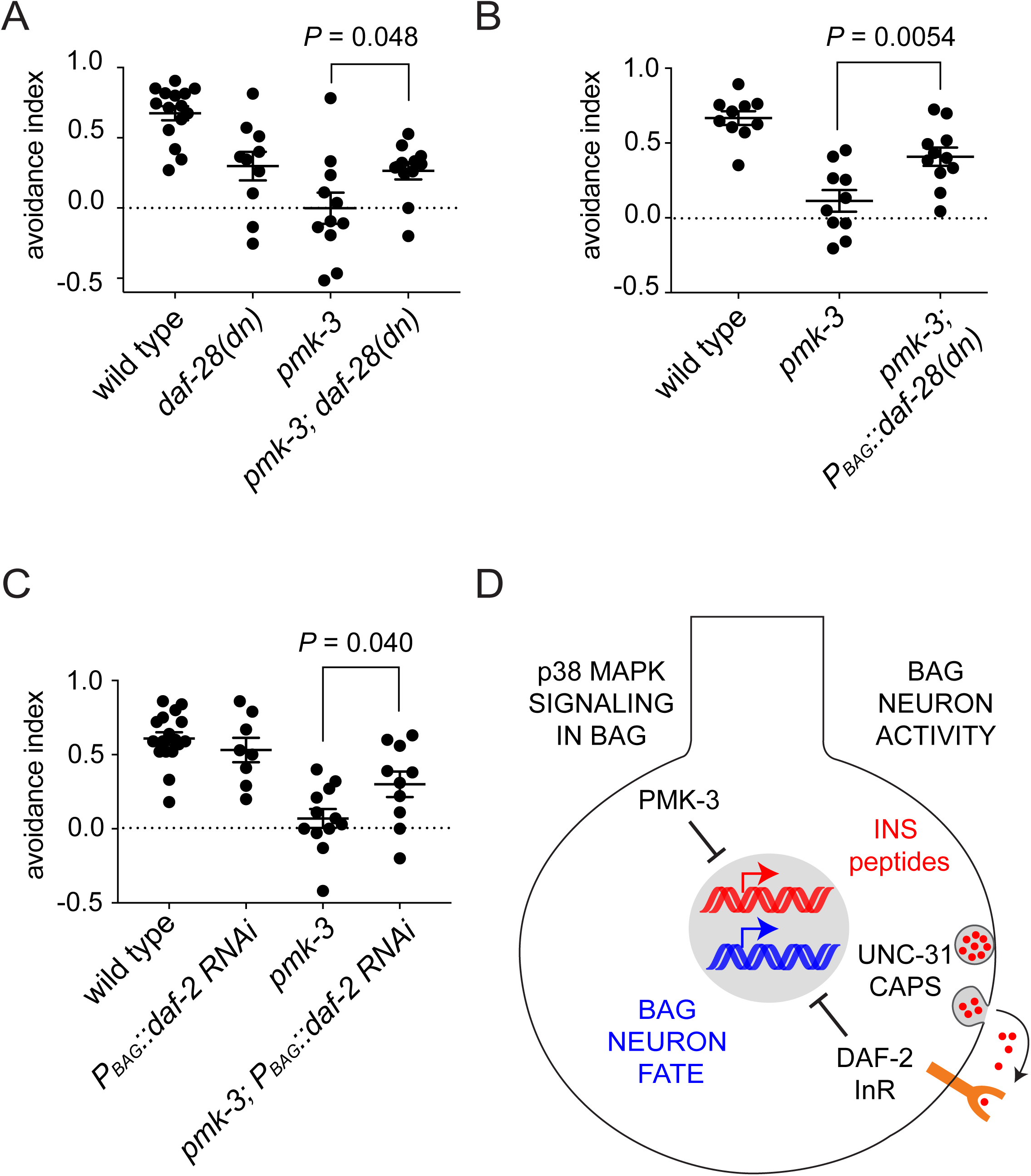
Disruption of autocrine insulin signaling restores function to *pmk-3* mutant BAG neurons. A. CO_2_-avoidance indices of the wild type, *daf-28(sa191), pmk-3(ok169)*, and *pmk-3(ok169); daf-28(sa191)* mutants. B. Reducing insulin secretion in *pmk-3(ok169)* mutant BAG cells by overexpressing *daf-28(sa191), P*_*BAG*_*∷daf-28(sa191)*, significantly restored CO_2_ avoidance behavior to *pmk-3(ok169)* mutants. C. BAG cell-targeted knockdown of the insulin receptor *daf-2* significantly restores CO_2_ avoidance behavior to *pmk-3(ok169)* mutant animals. N ≥ 10 trials. *P* values were determined by an unpaired t-test. D. Model for p38 MAPK regulation of BAG neuron development via regulation of an activity-dependent autocrine insulin signal.

## Discussion

We have found that the p38 MAP kinase PMK-3 controls development of *C. elegans* chemosensory BAG neurons through an unexpected mechanism: the regulation of an activity-dependent autocrine insulin signal (**Figure 7D**). This insulin signal is regulated at the level of gene transcription. *pmk-3* mutant sensory neurons overexpress transcripts encoding insulin-like peptides (ILPs). As a consequence, their BAG cells experience elevated DAF-2/Insulin Receptor (InR) signaling, which antagonizes expression of a neuropeptide gene essential for their function. We further show that the neuronal defects caused by loss of PMK-3 and concomitant increases in ILP production are abolished in mutants with defects in activity-dependent neuronal secretion. The autocrine insulin signal that controls expression of a chemosensory neuronal fate is, therefore, controlled both by a cell-intrinsic PMK-3-dependent genetic mechanism and by neural activity, which controls secretion of ILPs. This mechanism neatly integrates neural activity with a developmental program of gene expression to regulate a neuronal cell fate.

Remarkably, although ILP genes represent only a fraction of the genes that are dysregulated by loss of PMK-3 we found that it is possible to restore BAG-neuron function to *pmk-3* mutants by only manipulating insulin signaling in BAG cells. Because of the central role for ILP signaling in the etiology of the *pmk-3* mutant phenotype and because this mechanism converges on regulation of genes with defined roles in neuron-function, it will be of great interest to understand molecular mechanisms by which PMK-3 regulates ILP expression and how DAF-2/InR signaling regulates expression of FLP-17 neuropeptides.

### Activity-dependent autocrine growth-factor signaling in the nervous system

A key feature of ILP signaling in BAG neuron development is that BAG neurons themselves supply the ILPs that regulate their differentiation. Recently, a number of studies have revealed the functional importance of activity-dependent autocrine signals in circuit development (reviewed in (Herrmann and Broihier 2018)). In mammalian hippocampus, an activity-dependent autocrine Insulin-like Growth Factor 2 (IGF2) signal stabilizes synapses made by dentate granule cells onto their postsynaptic partners (Terauchi et al. 2016), and in visual cortex a subset of interneurons generates an activity-dependent autocrine IGF1 signal that regulates the strength of their inhibitory synaptic inputs (Mardinly et al. 2016). Non-insulin like growth factors also function as autocrine signals during nervous system development. Brain-Derived Neurotrophic Factor (BDNF) and Bone Morphogenic Protein (BMP) - like factor homolog have been identified as autocrine signals that regulate synapse development and function in mammalian hippocampus and at the insect neuromuscular junction, respectively (James et al. 2014; Harward et al. 2016).

Because these autocrine signals are released in an activity-dependent manner, they are able to trigger a mechanism that translates neuronal activity into intracellular signals to regulate gene expression. There are well studied mechanisms that couple neural activity to gene expression via intracellular calcium signaling *e.g.* CREB signaling (West and Greenberg 2011). Gene regulation by autocrine activity-regulated growth factor signals might, however, differ in functionally important ways. An autocrine signal might have a different threshold for activation by neural activity than CREB-dependent mechanisms, allowing for different patterns of neural activity to engage different gene regulatory mechanisms. Once engaged, autocrine signals might act on different timescales, and they likely regulate gene targets distinct from those regulated by CREB. Interestingly, some aspects of BAG neuron development are regulated by CREB (Rojo Romanos et al. 2017). Neural activity, therefore, controls the development of BAG neurons through multiple mechanisms, and these cells will be excellent models for studying the distinct roles in nervous system development of CREB-based excitation-transcription coupling and gene regulation by activity-regulated autocrine signals.

### Insulin signaling and nervous system development

During development of *C. elegans* BAG neurons, ILPs inhibit expression of a fully differentiated and functional BAG neuron fate. This role for ILPs differs from their established function as regulators of cell proliferation during nervous system development in both insects and mammals (Fernandez and Torres-Aleman 2012). During development of the *Drosophila* nervous system, ILPs are produced by glial cells in response to nutritive cues, and trigger adjacent neural stem cells to exit quiescence and begin dividing (Chell and Brand 2010; Sousa-Nunes et al. 2011). In mammals insulin-like signaling factors promote neural stem cell proliferation; changing IGF1 levels in the developing rodent brain correspondingly alters brain size (D’Ercole et al. 1996). Mammalian IGF-1 has been shown to function as a mitogenic signal during corticogenesis, and recruits neural progenitors to the cell-cycle (Mairet-Coello et al. 2009). A related factor - IGF2 - promotes the maintenance and expansion of neural stem cells in mammals (Ziegler et al. 2014), and also regulates the proliferation of cerebral cortical progenitors (Lehtinen et al. 2011). We observed a role for ILPs in nervous system development that is distinct from their known function as regulators of proliferation, and found that ILPs repress the differentiation of a specific neuron-type. Both functions of ILPs in nervous system development have in common, however, that they delay the appearance of fully differentiated and functional neurons – either by promoting proliferation of neuronal stem cells at the expense of their differentiation, or by inhibiting expression of specific neuronal cell fates.

*C. elegans* express a large number of ILPs, many in the nervous system. Insulin-like receptor and its associated signaling factors are also expressed throughout the *C. elegans* nervous system, as is PMK-3, which we show regulates ILP gene expression. It is, therefore, likely that the ILP-dependent mechanism that we have found in developing chemosensory neurons, also functions to regulate the differentiation of other neuron-types in *C. elegans*. Furthermore, the molecular constituents of ILP signaling pathways are conserved between nematodes and vertebrates. We suggest, therefore, that autocrine ILP signals might play similar roles in the regulation of neuronal cell fates during development of the vertebrate nervous system.

## Materials and Methods

### Strains

All strains used in this study were cultivated under standard conditions (Brenner 1974) at 20°C, and are listed in **Supplementary File 1**. The *daf-16(wz151)* deletion strain was generated by CRISPR/Cas9-mediated mutagenesis using Cas9 ribonucleoprotein (Paix 2015), along with a co-CRISPR strategy to increase efficiency (Kim et al. 2014), as previously described (Zamanian et al. 2018). Transgenic animals were generated via microinjection as previously described (Mello 1991).

### Plasmids

Plasmids used in this study were made using Gibson assembly (Gibson et al. 2009) and are listed in **Supplementary File 2**. The *daf-28(sa191)* cDNA was purchased as a gene block from Integrated DNA Technologies and cloned into the pPD49.26 fire vector using Gibson cloning.

### Microscopy

Animals were mounted on 2% agarose pads made in M9 medium and immobilized with 30 mM sodium azide (NaN_3_). Fluorescence and differential interference contrast (DIC) micrographs were acquired with a Zeiss Axioimager M2 upright microscope equipped with an EM-CCD camera (Andor) using a 100x objective (1.4 N.A.). Z-stacks were obtained with an LSM700 laser-scanning confocal microscope (Zeiss) using a 40x objective (1.4 N.A.). Maximum projections of image stacks were generated with Fiji (Schindelin et al. 2012).

### CO_2_ avoidance assays

CO_2_ avoidance assays were performed as previously described (Brandt and Ringstad 2015). Briefly, a total of 40-50 adult hermaphrodites were confined to a custom-made chamber on an unseeded 10-cm NGM plate fitted with inlets of air and 10% CO_2_. Gas mixes were pushed into the chamber at 1.5 mL/min using a syringe pump (New Era, Inc.). After 35 minutes, an avoidance index (A.I.) was computed according to the following equation: 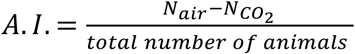.

### in vivo calcium imaging

*In vivo* calcium imaging was performed as previously described (Smith et al. 2013; Brandt and Ringstad 2015) using the ratiometric calcium indicator YC3.60 (Nagai et al. 2004).

### Quantification of GRASP puncta

The number and size of GRASP puncta were quantified using Fiji (Schindelin et al. 2012). Z-stacks of the BAG synaptic zone were thresholded and subjected to particle analysis, which automatically drew Region of Interests (ROIs) around puncta whose fluorescence intensity was above background. The number of puncta was defined as the number of particles found in the synaptic zone, and the area of each particle was measured to determine the size of the GRASP puncta. Each particle was manually inspected to confirm that it contained one GRASP puncta; if a particle encompassed multiple puncta - separate ROIs were manually drawn around the individual puncta.

### Genetic suppressor screen for regulators of p38 MAPK-dependent BAG cell development

*pmk-3* mutant animals carrying the BAG fate marker *ynIs64[P*_*flp-17*_*∷GFP]* were mutagenized with 47 mM ethyl methane sulfate (EMS) as described (Brenner 1974), and screened in the F2 generation for restored GFP expression on a Leica M165 FC fluorescence dissecting microscope. Approximately 10% of *pmk-3* mutants express *P*_*flp-17*_*∷GFP*, therefore mutants were identified as candidate suppressors of *pmk-3* if more than 50% of their F3 progeny had restored GFP expression.

### Mapping and cloning of suppressor alleles

Initial round of screening identified two non-complementing alleles, *wz75* and *wz76*. High-resolution-SNP-mapping of *wz75* was performed by crossing *wz75* mutants to the polymorphic strain CB4856 and identifying crossovers using restriction fragment polymorphisms (snip-SNPs) (Davis et al. 2005), which placed *wz75* in a 5 Mbp interval on LG IV. Whole-genome sequencing revealed that *wz75* and *wz76* mutants carry mutations in *unc-31*: *wz75* contains a G → A mutation predicted to change Trp971 to an Amber stop, and *wz76* contains a G → A mutation predicted to disrupt a splice-donor/acceptor site. Further screening and complementation tests identified 3 additional alleles of *unc-31: wz112, wz127*, and *wz130*. Whole genome sequencing of *wz112* and *wz130* revealed that *wz112* is a G → A missense mutation predicted to change Asp422 to Asn, and *wz130* is a G → A nonsense mutation predicted to change Trp1114 to an Opal stop. Sanger sequencing of *wz127* showed that it is a C → T mutation predicted to change Gln15 to an Amber stop.

### Transgene Expression Analysis

Animals were immobilized as described above. Reporter transgene expression was quantified using the 20x (.8 NA) objective on a Zeiss Axioimager M2 upright microscope equipped with an EM-CCD camera (Andor). To determine the penetrance of transgene expression, we measured the number of BAG cells expressing the reporter in each animal. As an alternative method, we counted the percentage of animals expressing the reporter on a Leica M165 FC fluorescence dissecting microscope. To determine the levels of reporter transgene expression, we measured the mean pixel values in a 30 or 50-pixel-radius circular ROI centered on the BAG soma using Fiji (Schindelin et al. 2012). For all experiments, data was collected over three days.

### Heat shock experiments

*pmk-3(ok169)* mutant animals carrying a *Phsp*_*16.41*_*∷irk-1* transgene were shifted twice to 37°C for a half hour, with a one hour recovery at 20°C in between. After heat shock, animals resumed growth at 20°C. Heat-shocked embryos and larvae were assayed for gene expression when they reached adulthood, while adults were assayed 24 hours after heat shock.

### Cell culture and FACS Sorting for RNA-Seq

Embryonic cell cultures were prepared from wild-type and *pmk-3(wz31)* mutant animals expressing the BAG-specific and *pmk-3* independent marker *wzIs113[P*_*gcy-9*_*∷GFP]*, as previously described (Christensen 2002; Zhang 2002). In brief, embryos were isolated from synchronized populations of hypochlorite treated hermaphrodites, and dissociated into single cells by chitinase treatment. Cells were resuspended in L-15 medium supplemented with 10% FBS (Sigma) and antibiotics, and passed through a 5 μm syringe filter (Millipore). Cells were plated onto poly-D-lysine coated single-well chambered cover glasses (Lab-Tek II) and incubated overnight at 25°C.

GFP-labeled BAG neurons were isolated approximately 24 hours after dissociation using Fluorescence Activated Cell Sorting (FACS); sorted cells were confirmed to be >90% GFP positive by direct inspection on a fluorescent microscope. Non-fluorescent, ‘non-BAG’, cells were also collected. Dead cells were marked with propidium-iodide and excluded from sorted cells. Sorting was performed on a FACSAria Ilu SORP cell sorter using a 70 μm nozzle. Cells were sorted directly into RNA Extraction Buffer (10,000 cells/100 μL buffer), and RNA was purified using the Arcturus PicoPure RNA Isolation Kit (Thermo Fisher). RNA integrity and concentration were evaluated using an Agilent Bioanalyzer. RNA samples had an RNA Integrity score of at least 8, indicating that all samples were of high quality. Two biological replicates were prepared from each cell type.

### RNA-Seq Analysis of wild type and pmk-3 mutant BAG neurons

cDNA libraries were prepared from RNA (1-2 ng) using the Ovation RNA-Seq System V2 (NuGEN), multiplexed, and sequenced as 100 base pair paired end reads using the HiSeq 2500 (Illumina). Reads were aligned to the *Caenorhabditis elegans* genome and transcriptome (Wormbase WS243) using the STAR software package (Dobin et al. 2013). Gene expression quantification was performed using HTSeq (Anders 2014). Differential expression analysis was done using the DeSeq2 package (Love et al. 2014) in R. Heatmaps were generated using gplots (Warnes 2016) and RColorBrewer (Neuwirth 2014) R software packages. Read coverage histograms were generated from genomic alignments (BAM files) using Integrative Genomics Viewer (Thorvaldsdottir et al. 2013).

### DAF-16∷GFP Localization

Z-stacks were collected from animals expressing *P*_*BAG*_*∷daf-16∷GFP* as described above, and image analysis was performed in Fiji (Schindelin et al. 2012). For each cell, the maximum sum projection of 5 stacks (∼1 μm/stack) was generated. ROIs were drawn around the nucleus and cell body in a summated projection image, and the total amount of DAF-16∷GFP fluorescence was measured in each region. The amount of fluorescence in the cytoplasm was defined as the total amount of DAF-16∷GFP fluorescence in the cell body minus the amount in the nucleus. We then computed the ratio of nuclear DAF-16∷GFP fluorescence to cytoplasmic DAF-16∷GFP fluorescence to monitor insulin activity.

### Statistical Analysis

Standard error of the mean (SEM) and *P*-values for statistical analyses were calculated using GraphPad Prism Software.

## Acknowledgments

This work was supported by the National Institute of Health (R35 GM122573 to N.R. and F31 NS100360 to L.B.H.). We thank Oliver Hobert (Columbia University) and Jane E. Hubbard (NYU School of Medicine) for reagents, Farzana Khan and Eugene Rudensky for isolating *unc-31* alleles from suppressor screens, Nechama Schneider for assisting with measurements of *flp-17* expression levels, and Igor Dolgalev for assisting with bioinformatic analysis. Flow cytometry assisted cell sorting was performed by the Cytometry and Cell Sorting Laboratory at NYU School of Medicine. Whole-genome sequencing and differential gene expression analysis was performed by the NYU School of Medicine Genome Technology Center. Some strains were provided by the CGC, which is funded by the NIH Office of Research Infrastructure Programs grant P40 OD010440, and the Mitani lab (Tokyo Women’s Medical University, Japan).

## Author Contributions

L.B.H. and N.R. designed and performed experiments, interpreted data, and wrote the manuscript. J.P.B. performed genetic suppressor screens and mapping experiments.

## Supplementary Files

**Supplementary File 1. *C. elegans* strains used in this study.**

A table of the *C. elegans* strains used in this study.

**Supplementary File 2. Plasmids used in this study.**

A table of the plasmids constructed in this study, and a table of the plasmids injected and associated concentration.

**Supplemental Figure S1:**
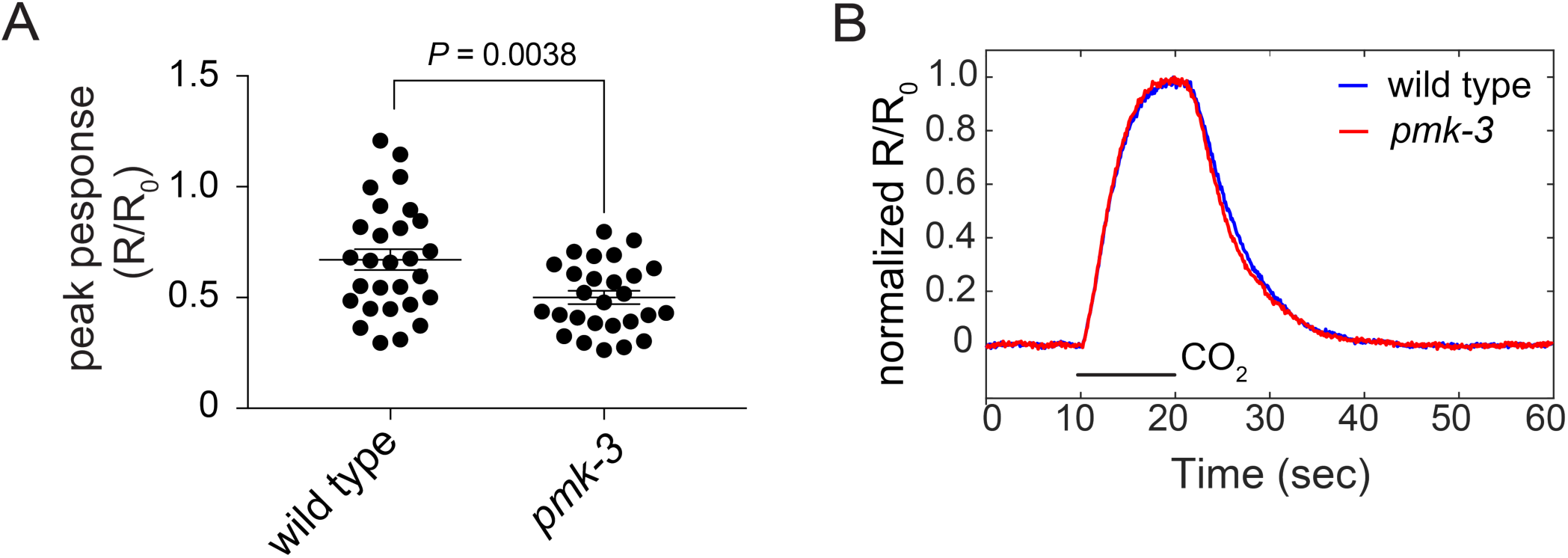
*pmk-3* mutant BAG neurons exhibit impaired chemotransduction in CO_2_-Sensing BAG neurons. A. Scatter plots showing the distribution of peak calcium responses (R/R_0_ values) of wild-type and *pmk-3(ok169)* mutant BAG neurons to 10% CO_2_ stimuli. *P* = 0.0038, unpaired t-test. N > 26 animals/genotype. Error bars represent SEM. B. Dynamics of average calcium responses of wild-type and *pmk-3(ok169)* mutant BAG neurons to 10% CO_2_ stimuli.

**Supplemental Figure S2:**
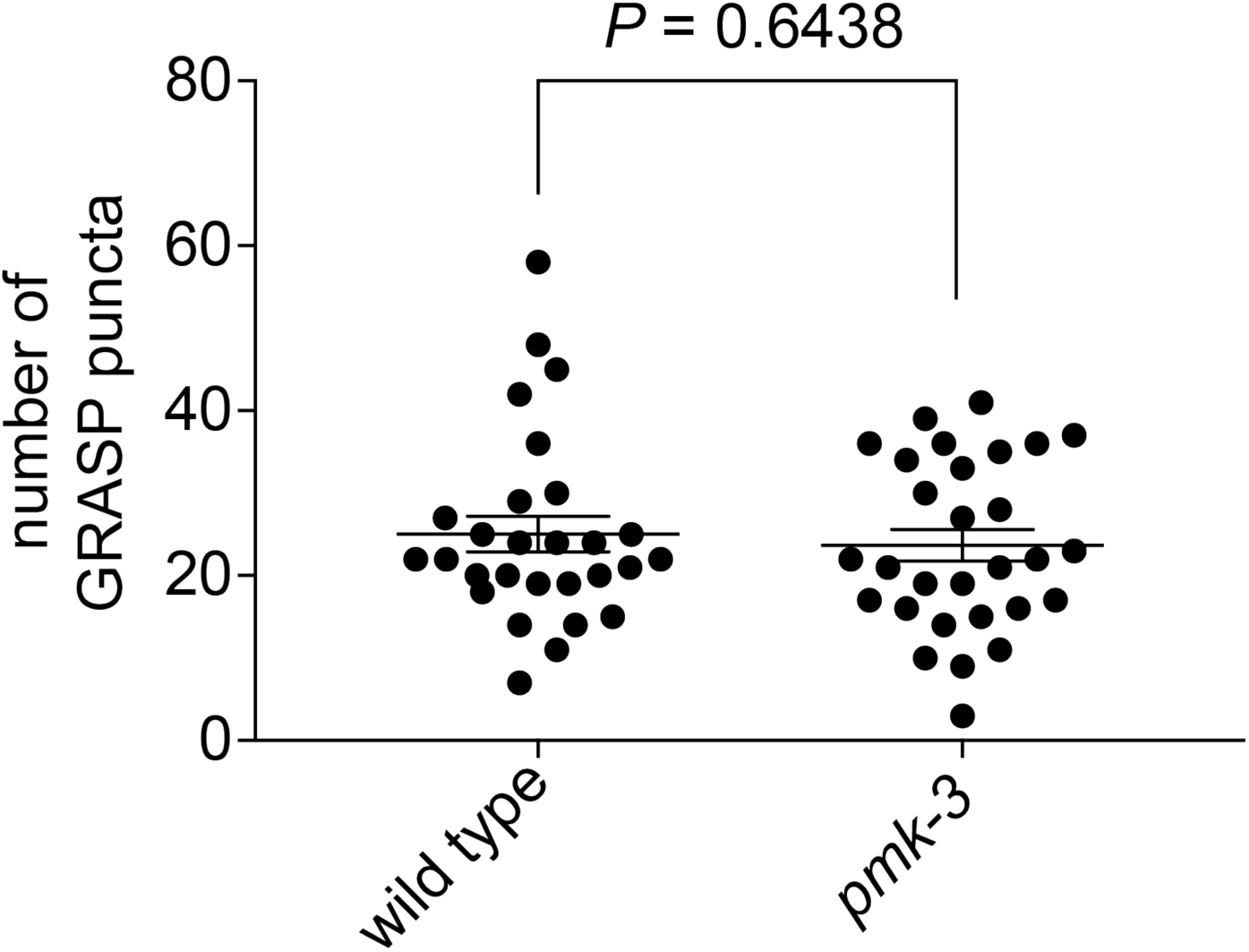
*pmk-3* mutant BAG neurons form normal number of synapses. Scatter plot showing the number of GRASP-puncta in wild-type and *pmk-3(ok169)* mutant BAG neurons. *P* = 0.6438, unpaired t-test. N > 28 animals/genotype. Error bars represent SEM.

**Supplemental Figure S3:**
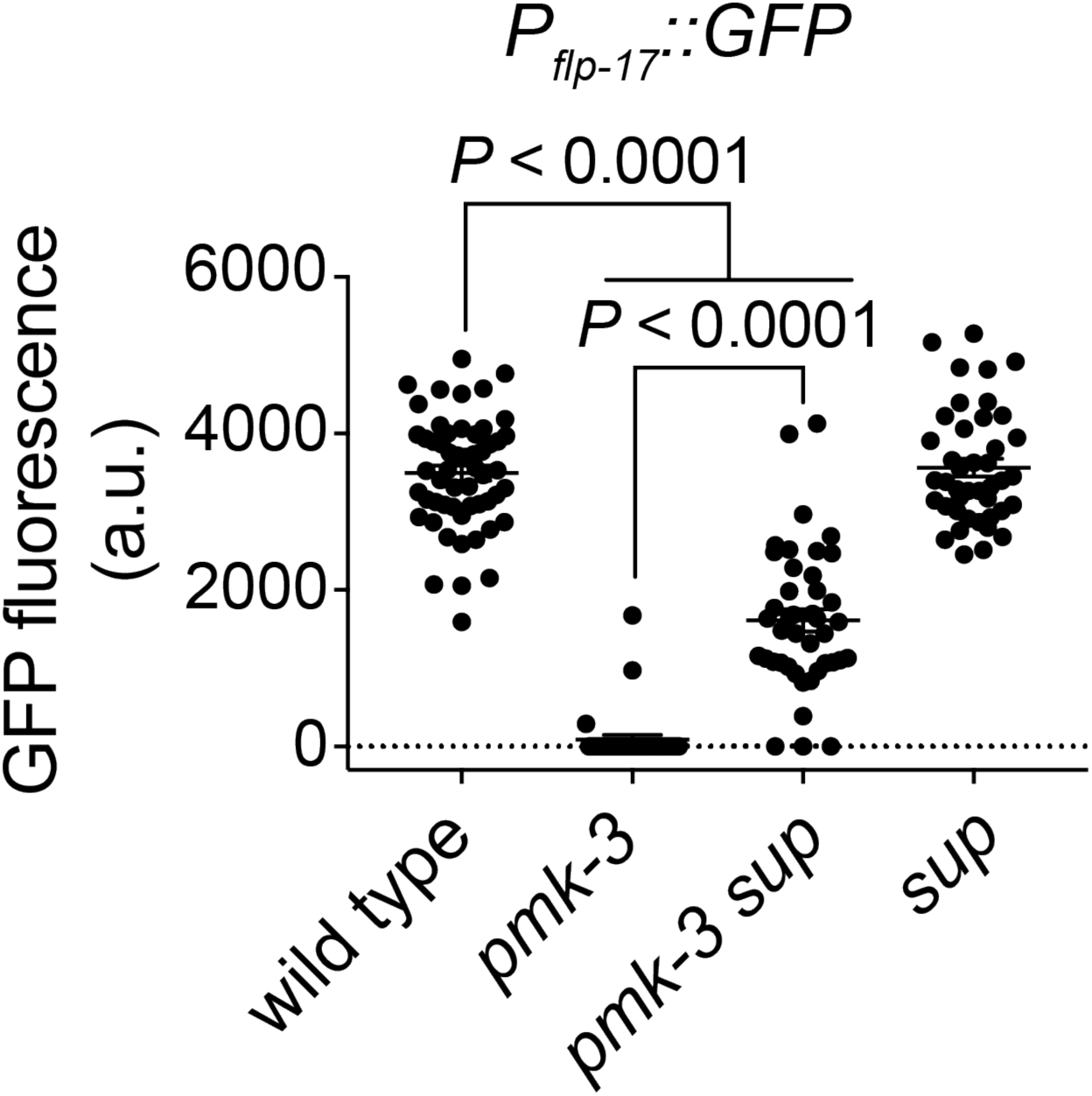
Suppressor mutations partially restore the levels of *flp-17* reporter expression to *pmk-3* mutant BAG cells. Levels of *P*_*flp-17*_*∷GFP* expression in the wild-type, *pmk-3(wz31), pmk-3(wz31) sup(wz75)*, and *sup(e169)* mutant animals (****) *P* < 0.0001, ordinary one-way ANOVA followed by Tukey’s multiple comparison test. N > 20 animals/genotype. (a.u.) arbitrary units. Error bars represent SEM.

**Supplemental Figure S4:**
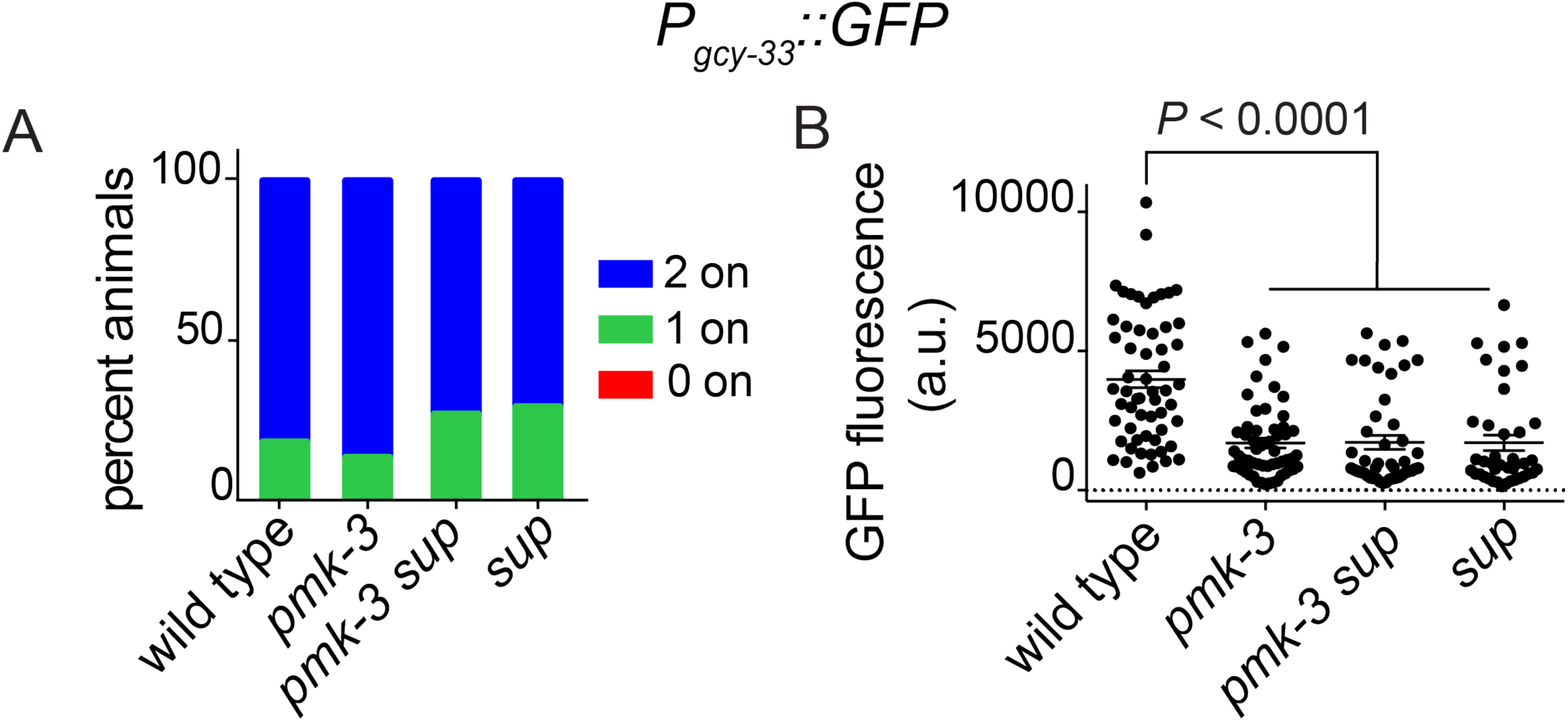
Suppressor regulates some, but not all, PMK-3 regulated genes in BAG neurons. (A). Frequency of *P*_*gcy-33*_*∷GFP* expression in the wild-type, *pmk-3(wz31), pmk-3(wz31) sup(wz75)*, and *sup(e169)* mutant animals. (B). Levels of *P*_*gcy-33*_*∷GFP* expression in the wild-type, *pmk-3(wz31), pmk-3(wz31) sup(wz75)*, and *sup(e169)* mutant animals. (****) *P* < 0.0001, ordinary one-way ANOVA followed by Tukey’s multiple comparison test. N > 20 animals/genotype. (a.u.) arbitrary units. Error bars represent SEM.

**Supplemental Figure S5:**
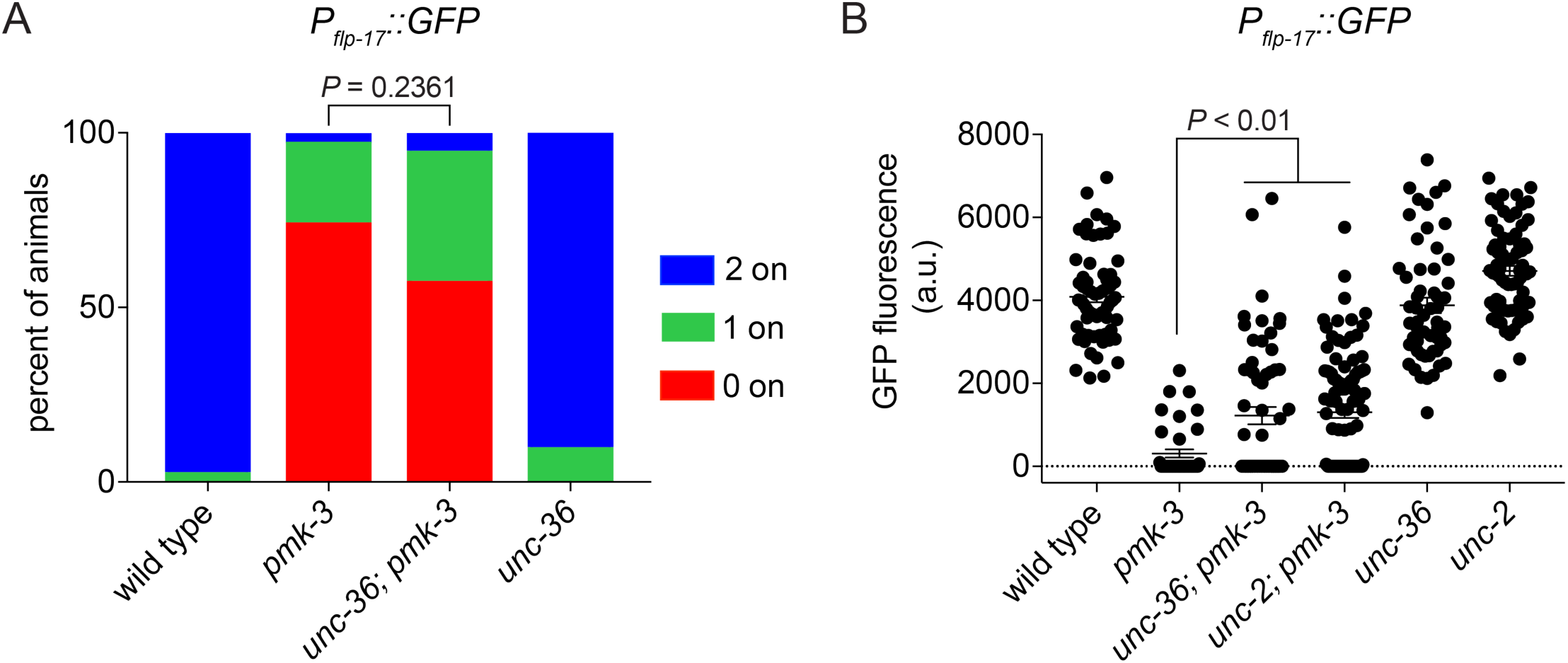
Mutations in NPQ-type Voltage Gated Calcium Channel subunits restore *flp-17* expression to *pmk-3* mutant BAG neurons. (A). Penetrance of *P*_*flp-17*_*∷GFP* expression in the wild-type, *pmk-3(ok169), unc-36(e251); pmk-3(ok169)*, and *unc-36(e251)* mutant animals. *P* = 0.2361, chi-square test. N ≥ 30 animals/genotype. (B). Levels of *P*_*flp-17*_*∷GFP* expression in the wild-type, *pmk-3(ok169), unc-36(e251); pmk-3(ok169), pmk-3(ok169); unc-2(e55), unc-36(e251)*, and *unc-2(e55)* mutant animals. *P* = 0.0028 for *pmk-3* vs *unc-36; pmk-3* and *P* = 0.0003 for *pmk-3* vs *pmk-3; unc-2*, ordinary one-way ANOVA followed by Tukey’s multiple comparison test. N ≥ 30 animals/genotype. (a.u.) arbitrary units. Error bars represent SEM.

**Supplemental Figure S6:**
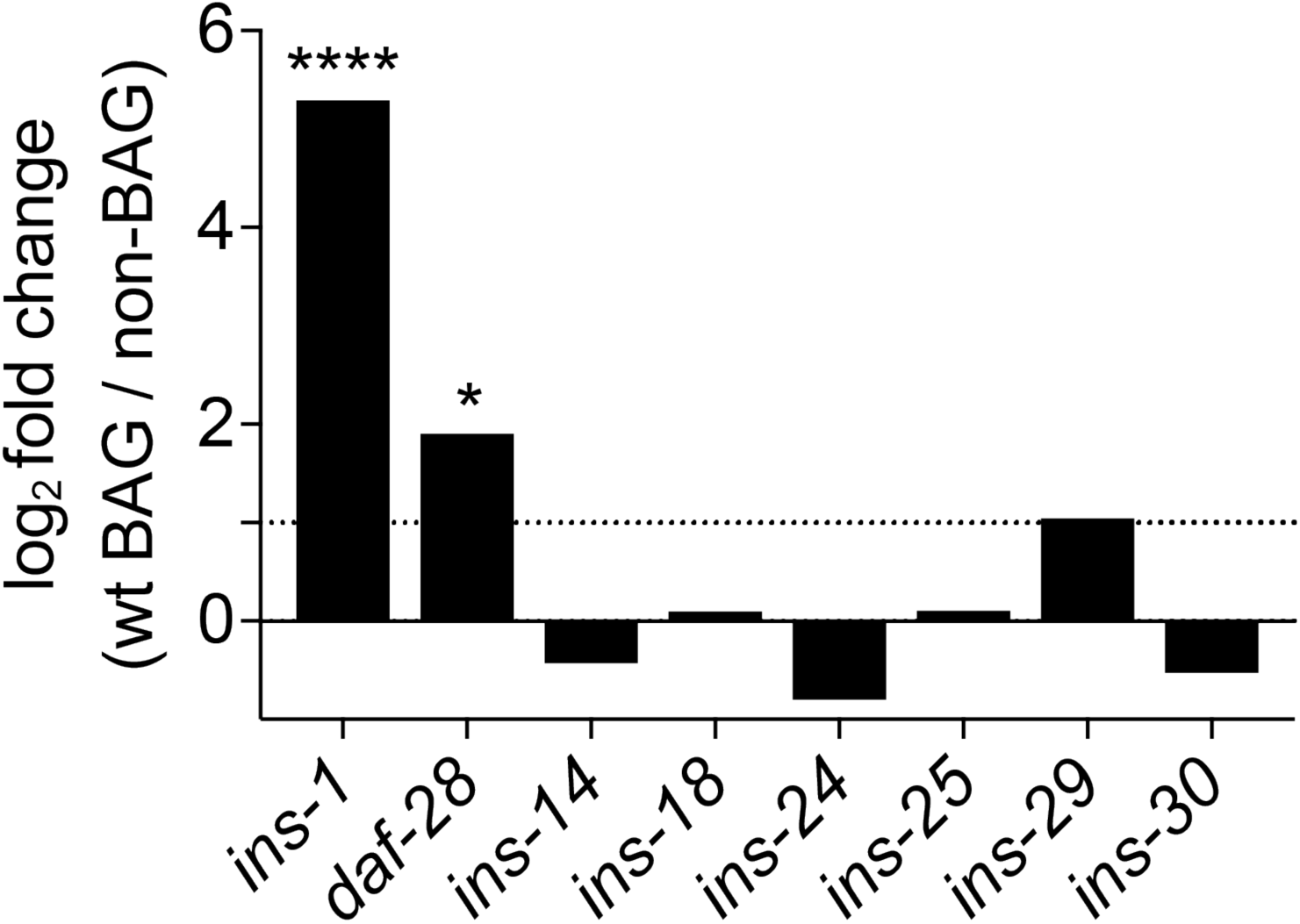
The insulin like peptides, *ins-1* and *daf-28*, are enriched in wild-type BAG cells. Fold changes of gene expression in wild type BAG cells versus non-BAG cells for *ins-*1 and ILPs that are enriched in *pmk-3* mutant BAG cells. (*) *P* < 0.05, (****) *P* < 0.0001.

**Supplemental Figure S7:**
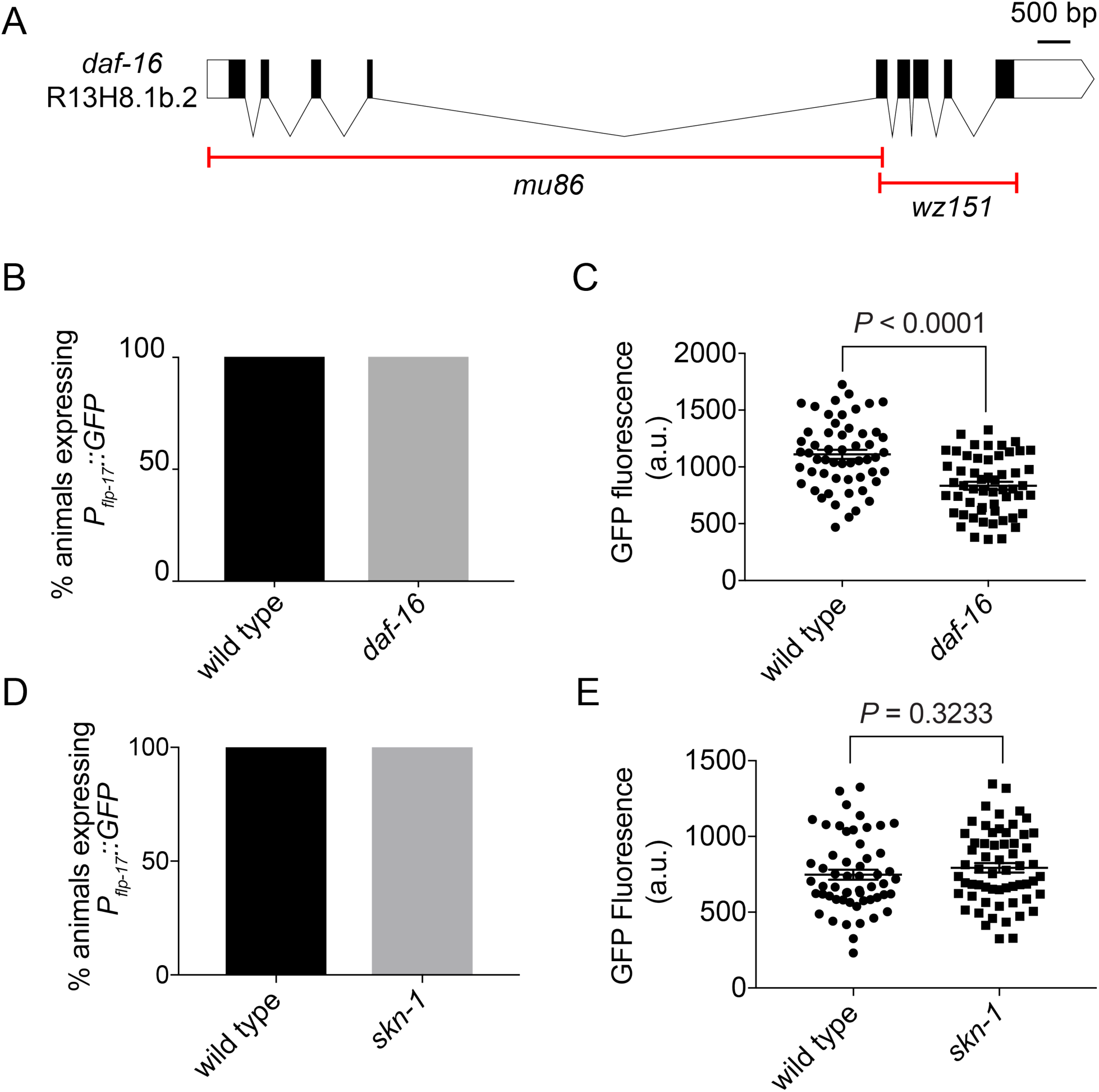
Loss of DAF-16 reduces levels of *flp-17* expression. (A). Structure of the *daf-16* genetic locus showing the canonical null allele, *mu86*, and the *wz151* 2065 base pair deletion allele generated using CRISPR/Cas9 mutagenesis. (B). Percentage of wild-type and *daf-16(wz151)* mutant animals expressing *P*_*flp-17*_*∷GFP*. (C). Levels of *P*_*flp-17*_*∷GFP* fluorescence in the wild-type and *daf-16(wz151)* mutant animals. *P* < 0.0001, unpaired t-test. (D). Percentage of wild-type and *skn-1(zu67)* mutant animals expressing *P*_*flp-17*_*∷GFP*. Because *skn-1(zu67)* is maternal effect lethal, *skn-1(zu67)* homozygous mutants were picked from heterozygous mothers carrying the *mIs11[myo-2∷GFP]* balancer chromosome for analysis. (E). Levels of *P*_*flp-17*_*∷GFP* fluorescence in the wild-type and *skn-1(zu67)* mutant animals. *P* = 0.3233, unpaired t-test. (a.u.) arbitrary units. N > 25 animals/genotype. Error bars represent SEM.

